# mTORC1 controls glycogen synthase kinase 3β nuclear localization and function

**DOI:** 10.1101/277657

**Authors:** Stephen J. Bautista, Ivan Boras, Adriano Vissa, Noa Mecica, Christopher M. Yip, Peter K. Kim, Costin N. Antonescu

## Abstract

Glycogen synthase kinase 3β (GSK3β) phosphorylates and regulates a wide range of substrates involved in diverse cellular functions. Some GSK3β substrates, such as c-myc and snail, are nuclear-resident transcription factors, suggesting possible control of GSK3β function by regulation of its nuclear localization. Inhibition of mechanistic target of rapamycin (mTORC1) led to partial redistribution of GSK3β from the cytosol to the nucleus, and GSK3β-dependent reduction of the expression of c-myc and snail. mTORC1 is controlled by metabolic cues, such as by AMP-activated protein kinase (AMPK) or amino acid abundance. Indeed AMPK activation or amino acid deprivation promoted GSK3β nuclear localization in an mTORC1-dependent manner. GSK3β was detected in several distinct endomembrane compartments, including lysosomes. Consistently, disruption of late endosomes/lysosomes through perturbation of Rab7 resulted in loss of GSK3β from lysosomes, and enhanced GSK3β nuclear localization as well as GSK3β-dependent reduction of c-myc levels. This indicates that GSK3β nuclear localization and function is suppressed by mTORC1, and suggests a new link between metabolic conditions sensed by mTORC1 and GSK3β-dependent regulation of transcriptional networks controlling biomass production.

**Summary statement (15-30 words):** GSK3β nuclear localization and function is negatively regulated by the metabolic and mitogenic sensor mTORC1. mTORC1 control of GSK3β localization requires Rab7 and lysosomal membrane traffic.

## Introduction

Glycogen synthase kinase 3β (GSK3β) is a serine/threonine protein kinase that controls numerous aspects of cellular physiology such as proliferation, metabolism, and apoptosis (Beurel et al., 2015; Cormier and Woodgett, 2017; Doble and Woodgett, 2003; Sutherland, 2011). Dysregulation of GSK3β has been linked to various diseases such as insulin resistance/diabetes, Alzheimer’s disease and cancer (Jope and Johnson, 2004). GSK3β phosphorylates over 100 substrates, more than the typical number of substrates for most kinases (Beurel et al., 2015; Linding et al., 2007; Sutherland, 2011), thus illustrating the broad capabilities for control of cell physiology by GSK3β. Notably, GSK3β is further distinguished from other kinases by being basally active (Doble and Woodgett, 2003). Hence, many mechanisms likely exist to regulate GSK3β.

GSK3β activity is indeed regulated by phosphorylation on S9, mediated by kinases such as Akt, protein kinase C (PKC), and p90RSK, resulting in negative regulation of GSK3b activity (Cross et al., 1995; Delcommenne et al., 1998; Fang et al., 2000; Stambolic and Woodgett, 1994; Sutherland et al., 1993; Tsujio et al., 2000). Phosphorylation of other sites on GSK3b may also suppress GSK3β activity, such as that of S389 by p38 MAPK (Thornton et al., 2008). In addition to GSK3β phosphorylation, control of GSK3β action may be achieved by localization of GSK3β or some of its substrates into distinct cellular compartments, such as the nucleus, such that GSK3β may have limited and regulated access to certain substrates (Bechard and Dalton, 2009; Meares and Jope, 2007; Sutherland, 2011).

Several GSK3β substrates are transcription factors (Sutherland, 2011) localized largely to the nucleus, including c-myc (Gregory et al., 2003), snail (Zhou et al., 2004), C/EBPα and β (Ross et al., 1999; Tang et al., 2005), and CREB (Fiol et al., 1994). C-myc controls genes important for proliferation, metabolism and biomass production, and stem-cell self renewal (reviewed by (Dang, 2012; Dang et al., 2009; Kalkat et al., 2017)). Moreover, c-myc is an oncogene altered in many cancers (Kalkat et al., 2017), highlighting the need for precise regulation of its function. C-myc protein levels are controlled by GSK3β-dependent phosphorylation of T58 on c-myc (Gregory et al., 2003), leading to ubiquitin-dependent proteosomal degradation (Thomas and Tansey, 2011). Control of phosphorylation and/or degradation of these nuclear substrates by GSK3β may involve modulation of GSK3β nuclear localization. However, the identity of the cellular compartments in which GSK3β is localized, and how it moves from various cellular compartments to the nucleus is not well defined.

GSK3β localizes in part to membrane compartments in the cytoplasm. It is recruited to the plasma membrane via association with Axin (Zeng et al., 2008), impacting Wnt signaling to β-catenin (Wu and Pan, 2010). GSK3β is also detected on APPL1 early endosomes (Schenck et al., 2008). APPL1 acts as an adaptor protein to recruit Akt, facilitating GSK3β phosphorylation and inactivation on these early endosomes, thus impacting clathrin-mediated endocytosis (Reis et al., 2015) and cell survival (Schenck et al., 2008). GSK3β also localizes to lysosomes (Li et al., 2016) and controls lysosomal acidification (Azoulay-Alfaguter et al., 2015). Hence, GSK3β may localize to multiple distinct endomembrane compartments including the plasma membrane, early endosomes and lysosomes, with distinct functions at each locale.

GSK3β exhibits nuclear localization under certain conditions including in response to apoptotic signals induced by heat shock or staurosporine treatment (Bijur and Jope, 2001), S-phase of the cell cycle (Diehl et al., 1998), replicative senescence in fibroblasts (Zmijewski and Jope, 2004), and loss of phosphatidylinositol-3-kinase (PI3K)-Akt signaling in embryonic stem cells (Bechard and Dalton, 2009). Site-directed mutagenesis studies revealed that nuclear localization of GSK3β requires a bipartite nuclear localization sequence (NLS) contained within resides 85-103 on GSK3β, and that nuclear localization was also modulated by the N-terminal 9 amino acids on GSK3β (Meares and Jope, 2007).

Collectively, these observations raise the question of whether the control of GSK3β nucleocytoplasmic shuttling could be an important mechanism to control its function by modulating access to nuclear substrates. Indeed nuclear localization of GSK3β induced by inhibition of PI3K or Akt leads to GSK3β-dependent phosphorylation of c-myc, leading to its degradation (Bechard and Dalton, 2009). However, it is not clear how PI3K-Akt signals impact GSK3β localization. Since this N-terminal region contains the S9 phosphorylation site, it is possible that Akt or other kinases capable of phosphorylation of this residue impact nuclear localization of GSK3β, although a GSK3β mutant (S9A) did not show obvious differences in nuclear localization (Meares and Jope, 2007).

While Akt may control GSK3β localization via direct phosphorylation of GSK3β on S9, this may be indirect and result from Akt-dependent activation of other signals, downstream of Akt, such as the mechanistic target of rapamycin complex 1 (mTORC1). Mitogenic activation of PI3K-Akt signals leads to inhibition of the Tuberous Sclerosis Complex 1/2 (TSC 1/2) (Inoki et al., 2002), activation of the GTPase Rheb (Inoki et al., 2003), and thus mTORC1 activation (Long et al., 2005; Tee et al., 2003). mTORC1 in turn controls many processes including metabolism, protein synthesis, cell growth and autophagy (recently reviewed by (Saxton and Sabatini, 2017)). In addition to mitogenic control, mTORC1 is also strongly controlled by metabolic cues. Amino acid levels are sensed by a mechanism involving the lysosomal V-ATPase and other sensors, leading to the recruitment and activation of mTORC1 at the lysosome under conditions of amino acid sufficiency (Bar-Peled et al., 2012; Zoncu et al., 2011). Further, activation of AMP-activated protein kinase (AMPK) by energy insufficiency, resulting from an increase in the cellular levels of AMP and ADP relative to ATP, leads to phosphorylation and activation of TSC2, thus suppressing mTORC1 (Inoki et al., 2006; Shaw et al., 2004). Hence, mitogenic and metabolic signals control mTORC1 activation.

While it is not known if PI3K-Akt signaling regulates GSK3β nuclear localization *via* engagement of mTORC1, several studies have reported that GSK3β enhances mTORC1 activity. GSK3β phosphorylates the mTORC1 subunit Raptor (Stretton et al., 2015), resulting in enhanced mTORC1 activity (Azoulay-Alfaguter et al., 2015; Stretton et al., 2015). GSK3β also negatively regulates mTORC1 signaling by binding (Ka et al., 2014) and phosphorylation of TSC2 (Inoki et al., 2006). Moreover, GSK3β binds to and regulates AMPK (Suzuki et al., 2013). Hence, GSK3β controls the mTORC1 and AMPK metabolic and mitogenic sensors. However, the possibility of a reciprocal regulation of GSK3β by signals from mTORC1 and AMPK, impacting GSK3β nuclear localization and thus access to substrates therein such as c-myc, has so far been unexamined.

Here, we examine mTORC1 regulation of GSK3β nuclear localization and function. To do so, we use pharmacological and other approaches to manipulate mitogenic or metabolic signals and examine GSK3β localization to various endomembrane compartments and nucleus as well as GSK3β-dependent functions associated with nuclear GSK3β. We find a novel regulatory axis sensing mitogenic signals, metabolic cues and membrane traffic at the late endosome/lysosome that modulates GSK3β nuclear localization and function.

## Results

The PI3K-Akt signaling pathway controls GSK3β nucleocytoplasmic shuttling and thus access of GSK3β to nuclear targets either directly or via activation of the downstream kinase mTORC1. mTORC1 integrates both mitogenic (PI3K-Akt) and metabolic cues, and is localized to the lysosome once activated. Using a variety of strategies to manipulate mitogenic and metabolic signals converging on mTORC1 and lysosomal membrane traffic, we examined how mTORC1 regulates GSK3β nuclear access and function.

### mTORC1 controls GSK3 β nuclear localization and c-myc expression

To determine whether mTORC1 regulates GSK3β localization and function downstream of PI3K/Akt, we first examined the effect of the mTORC1 inhibitor rapamycin on c-myc expression. Treatment of ARPE-19 cells (RPE henceforth) with 1 μg/mL rapamycin caused a time-dependent decrease in c-myc expression, reaching 57 ± 4.8% after 2 hours of rapamycin treatment (n = 6, p < 0.05, **Fig. 1A**). Importantly, co-treatment with 10 μM of the GSK3β kinase inhibitor CHIR99021 blunted the decrease in c-myc expression elicited by rapamycin treatment (**Fig. 1A**). Consistent with this result, rapamycin treatment also elicited a reduction in expression of the transcription factor snail, an effect also blunted by co-treatment with CHIR 99021 (**Fig. 1B**).

**Figure 1.**
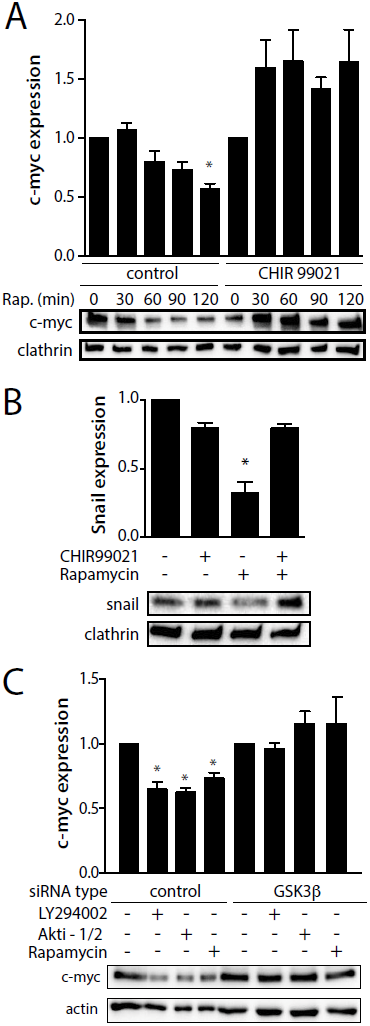
mTORC1 inhibition decreases c-myc and snail expression in a GSK3β -dependent manner. (***A-B***) RPE cells were treated with 1 μM Rapamycin, in the presence or absence of 10 μM CHIR 99021 for the indicated times (A) or 1 h (*B*). Shown are representative immunoblots of whole-cell lysates probed with anti-c-myc (A), anti-snail (B) or anti-clathrin heavy chain (load) antibodies. Also shown are mean c-myc levels ± SE, n = 6, * *p* < 0.05 (A), or mean snail levels n=3; *, p < 0.05 (B), relative to that in the control conditions (no inhibitor treatment). (***C***) RPE cells were transfected with siRNA targeting GSK3β or non-targeting siRNA (control), then treated with either 10 μM LY294002, 5 μM Akti-1/2, or 1 μM Rapamycin for 1 h. Shown are representative immunoblots of whole-cell lysates probed with anti-c-myc or anti-actin (load) antibodies, as well as mean c-myc levels, n = 4; * p < 0.05, relative to that in the control conditions (no inhibitor treatment).

We next used siRNA gene silencing of GSK3β, which resulted in a 91 ± 4.7% reduction of GSK3β protein levels (n = 3, p < 0.05, **Fig. S1A**). While RPE cells also express the paralog GSK3α, silencing of GSK3β was specific and did not impact expression of GSK3α (**Fig. S1B**). Cells subjected to silencing of GSK3β exhibited no change in c-myc upon inhibition of sequential signals in the PI3K-Akt-mTORC1 axis, achieved by treatment with either LY294002, Akti-1/2, or rapamycin, respectively (**Fig. 1C**). In contrast, each inhibitor effectively reduced c-myc expression in cells subjected to non-targeting (control) siRNA treatment (**Fig. 1C**). Taken together, these results indicate that PI3K-Akt signals converge on mTORC1 to enhance c-myc levels in a manner that requires the regulation of GSK3β.

To determine how PI3K-Akt-mTORC1 signals control c-myc expression in a GSK3β-dependent manner, we examined the localization and levels of endogenous GSK3β and c-myc. Consistent with a previous report (Bechard and Dalton, 2009), in cells grown in serum with an active PI3K-Akt-mTORC1 axis, GSK3β primarily localizes within the cytosol and appears mostly excluded from the nucleus (**Fig. 2A**). We confirmed the specificity of detection of endogenous GSK3β by immunofluorescence microscopy following GSK3β silencing (**Fig. S1C**). In contrast, and as expected (Abrams et al., 1982; Hann et al., 1983; Smith et al., 2004), c-myc localizes virtually entirely within the nucleus under these conditions (**Fig. 2A**). Thus, under conditions in which mTORC1 is active, GSK3β and c-myc are compartmentalized separately within the cytosol and nucleus, respectively.

**Figure 2.**
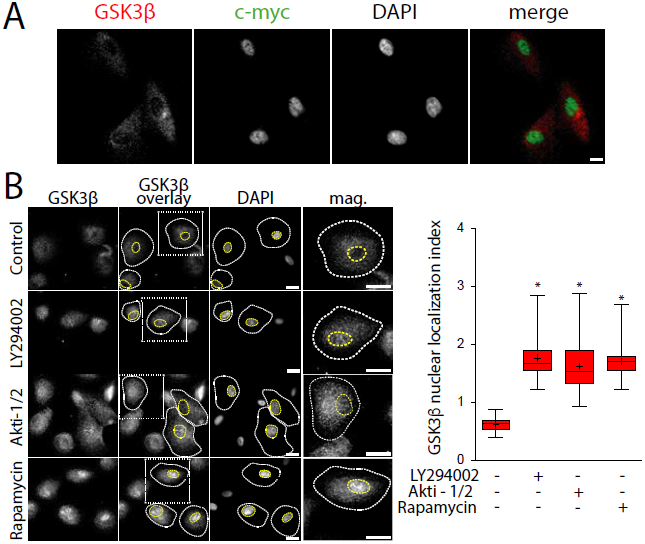
Inhibition of PI3K/Akt/mTORC1 signals promotes GSK3β nuclear localization. (***A***) Representative images obtained by widefield epifluorescence microscopy of control RPE cells (no inhibitor treatment) stained to detect endogenous GSK3β or c-myc, with DAPI stain to identify the nucleus, scale = 20 μm. (***B***) RPE cells were treated with either 10 μM LY294002, 5 μM Akti-1/2, or 1 μM Rapamycin for 1 h, then fixed and stained to detect endogenous GSK3β. Shown (left panel) are micrographs obtained by widefield epifluorescence microscopy representative of 3 independent experiments, scale = 20 μm. Also shown for each condition as ‘GSK3β overlay’ are sample cellular and nuclear outlines, and a box corresponding to a magnified image of a single cell. Also shown (right panel) is the mean GSK3β nuclear localization index ± SE (n = 3, >30 cells per condition per experiment); *, *p* < 0.05 relative to control conditions (no inhibitor treatment).

We next determined how PI3K-Akt-mTORC1 signaling regulates GSK3β localization. Treatment of RPE cells with either LY294002, Akti-1/2, or rapamycin to perturb PI3K, Akt or mTORC1, respectively resulted in robust and significant (n = 3, p < 0.05) increase in nuclear GSK3β, measured by the ratio of nuclear to cytosolic mean fluorescence intensities of GSK3β which we term the GSK3β nuclear localization index (**Fig. 2B**). Importantly, the effect of rapamycin treatment on GSK3β nuclear translocation and snail protein levels was also observed in MDA-MB-231 breast cancer cells (**Fig. S1D-E**), demonstrating that the mTORC1-dependent control of GSK3β is not unique to RPE cells. Furthermore, inhibition of the PI3K-Akt-mTORC1 axis also resulted in robust nuclear localization of GSK3α (**Fig. S1E**), a paralog of GSK3β with highly similar kinase domains but unique terminal motifs (Cormier and Woodgett, 2017; Woodgett, 1990). These results indicate that PI3K-Akt signals act via control of mTORC1 to regulate GSK3β nuclear localization, as well as that of GSK3α.

To test the importance of Ran in mTORC1-dependent GSK3β nuclear translocation, we examined the impact of Ran GTP-binding mutants on GSK3β localization. We expressed wild-type (WT) Ran or one of two Ran mutants, Ran T24N and G19V, which are constitutively GDP-or GTP-bound, respectively (Carey et al., 1996). Cells expressing WT Ran exhibited little nuclear GSK3β in the control condition, but a robust localization of GSK3β in the nucleus was observed upon treatment with rapamycin (**Fig. 3**, upper panels, and quantification, lower panel). In contrast, cells expressing Ran T24N exhibited nuclear GSK3β in both control and rapamycin-treated conditions (**Fig. 3**), consistent with Ran-GDP acting to facilitate nuclear import (Carey et al., 1996). Further, cells expressing Ran G19V exhibited mostly cytosolic GSK3β in both control and rapamycin-treated conditions, consistent with this mutant blocking Ran-dependent nuclear import (**Fig. 3**). These results indicate that GSK3β undergoes Ran-dependent nucleocytoplasmic shuttling and Ran-dependent nuclear import that is regulated by mTORC1.

**Figure 3.**
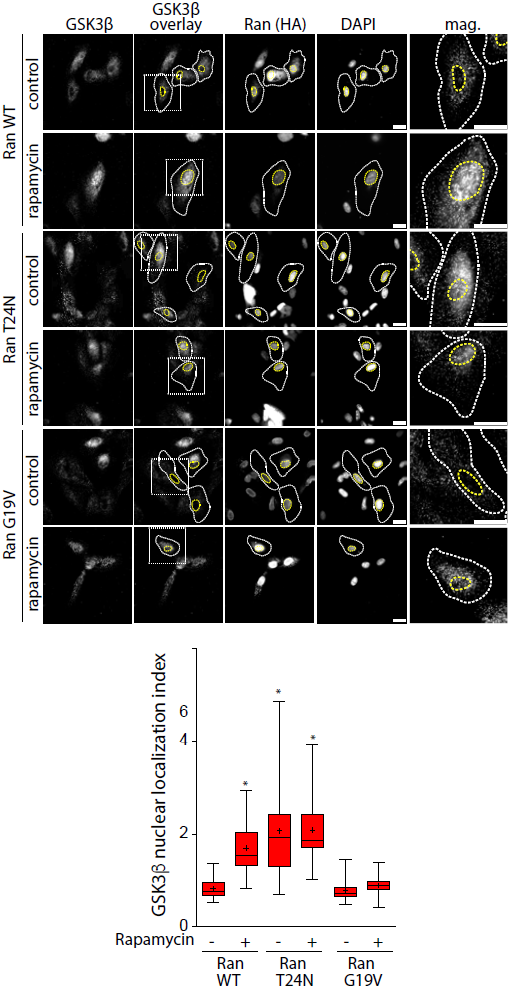
Rapamycin-induced GSK3β nuclear localization is Ran-dependent. (***A***) RPE cells were transfected with plasmids encoding HA-tagged wild-type (WT), T24N or G19V Ran, then treated with 1 μM Rapamycin for 1 h, followed by detection of endogenous GSK3β and exogenous HA-tagged Ran proteins. Shown (top panel) are micrographs obtained by widefield epifluorescence microscopy representative of 3 independent experiments, scale = 20 μm. Also shown for each condition as ‘GSK3β overlay’ are sample cellular and nuclear outlines, and a box corresponding to a magnified image of a single cell. Also shown (bottom panel) is the mean GSK3β nuclear localization index ± SE (n = 3, >30 cells per condition per experiment); *, *p* < 0.05 relative to control conditions (no inhibitor treatment).

### Metabolic cues regulate GSK3 β nuclear localization via mTORC1

As mTORC1 is regulated by both mitogenic (PI3K-Akt) signals as well as metabolic cues, we next examined how each of these signals contributes to the control of GSK3β nuclear localization. AMPK is activated via ATP insufficiency, and negatively regulates mTORC1 signaling through phosphorylation and activation of TSC2 (Inoki et al., 2006; Shaw et al., 2004). Consistent with the effects of mTORC1 inhibition by rapamycin, treatment with the AMPK activator A769662 resulted in robust GSK3β nuclear localization (**Fig. 4A**). Importantly, AMPK and mTORC1 exhibit reciprocal negative regulation (Inoki et al., 2012). As such, GSK3β nuclear localization could conceivably be the direct result of loss of mTORC1 signals, or an increase in AMPK activation, both of which would be expected to occur upon treatment with either rapamycin or A769662. To dissect a role for mTORC1 versus AMPK in control of GSK3β nuclear localization, we used the AMPK inhibitor compound C (Ross et al., 2015). Cells treated with compound C exhibited a rapamycin-dependent increase in GSK3β nuclear localization comparable to that observed in cells treated with rapamycin but not compound C (**Fig. 4A**). This indicates that AMPK activity is dispensable for GSK3β nuclear localization induced by mTORC1 inhibition. As GSK3β forms a complex with AMPK (Suzuki et al., 2013), we also tested whether AMPK may have a kinase-independent, structural role in regulation of GSK3β. However, silencing of AMPK did not impact GSK3β nuclear localization (**Fig. S2**). Collectively, these results indicate that while AMPK activation also triggers an accumulation of nuclear GSK3β, this occurs as a result of AMPK-dependent inhibition of mTORC1 signals, and not as a result of direct action of AMPK on GSK3β nuclear localization.

**Figure 4.**
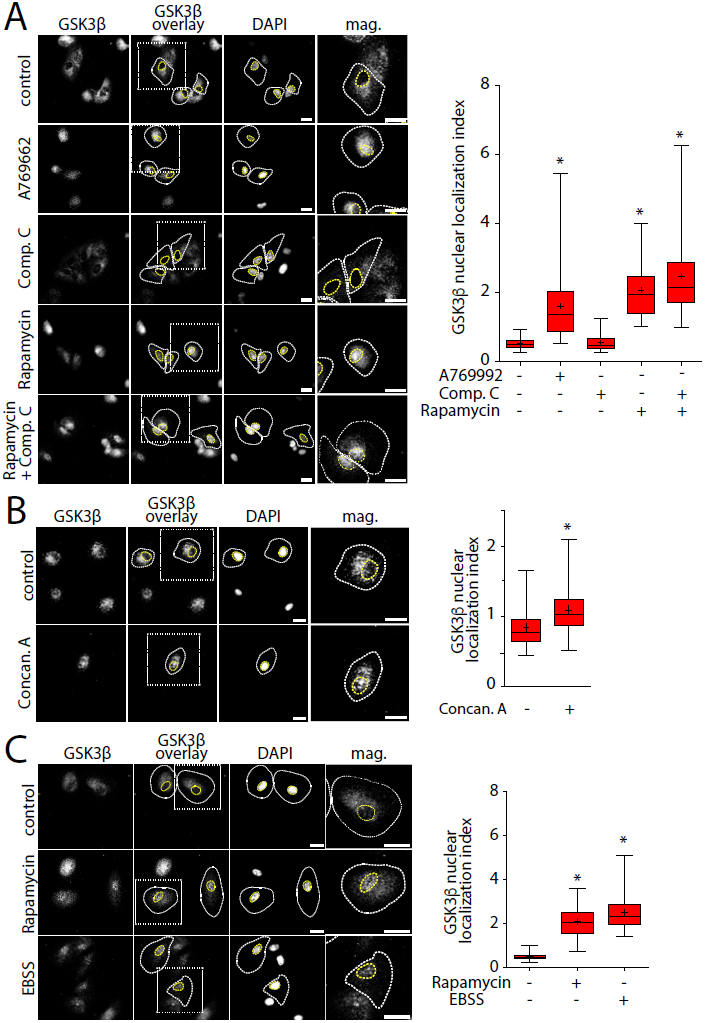
mTORC1 integrates multiple signals to control GSK3β nuclear localization. RPE cells were treated with either 100 μM A769662, 5 μM Compound C, or 1 μM Rapamycin, alone or in combination for 1h (***A***), or 1 μM Concanamycin for 1h (***B***), or incubated in amino acid-free EBSS media for 2h (***C***). Shown (left panels) in each case are micrographs obtained by widefield epifluorescence microscopy representative of 3 independent experiments, scale = 20 μm. Also shown for each condition as ‘GSK3β overlay’ are sample cellular and nuclear outlines, and a box corresponding to a magnified image of a single cell. Also shown (right panels) are the mean GSK3β nuclear localization indices ± SE (n = 3, >30 cells per condition per experiment); *, *p* < 0.05 relative to control conditions (no inhibitor treatment).

mTORC1 is activated by abundance of amino acids in a manner that requires the V-ATPase (Zoncu et al., 2011). To determine how amino acid-dependent activation of mTORC1 impacted control of GSK3β localization, we treated cells with the V-ATPase inhibitor Concanamycin A. Cells treated with Concanamycin A exhibited a significant enhancement of nuclear GSK3β relative to control (**Fig. 4B**). Consistent with this result, amino acid deprivation achieved via treatment of cells in amino acid depleted media (EBSS) also mimicked the effect of rapamycin treatment in RPE (**Fig. 4C**) as well as MDA-MB-231 (**Fig. S1C**) cells. These results indicate that amino acid sensing by mTORC1 contributes to the regulation of GSK3β nuclear localization.

mTORC1 inhibition also leads to induction of autophagy (Jung et al., 2009). We therefore tested whether autophagy is required for GSK3β nuclear localization upon mTORC1 inhibition with rapamycin. To inhibit autophagy induction, we treated cells siRNA targeting endogenous ULK (Saric et al., 2016), which resulted in a robust 77% ± 6.2 reduction of ULK expression (n = 3, p < 0.05, **Fig. S3A**). Cells treated with siRNA to silence ULK1 exhibited cytosolic GSK3β, which relocalized to the nucleus upon rapamycin treatment in a manner indistinguishable from cells treated with non-targeting siRNA (**Fig. S3B**). As autophagy induction has also been reported to lead to c-myc degradation (Cianfanelli et al., 2014), we also tested the effect of ULK1 silencing on rapamycin-induced c-myc expression. Surprisingly, silencing of ULK1 on its own reduced c-myc expression (**Fig. 3C**). Moreover, and in contrast to the findings of a previous study (Cianfanelli et al., 2014), impairment of autophagy induction by ULK1 silencing did not prevent the rapamycin-induced reduction in c-myc expression (**Fig. S3C**). Thus, GSK3β nuclear translocation and c-myc degradation observed upon mTORC1 inhibition are largely independent of autophagy induction. Instead, c-myc degradation upon mTORC1 inhibition is mediated by regulation of GSK3β nuclear localization and function.

### Control of GSK3β nuclear localization does not require GSK3β S9 phosphorylation

Akt phosphorylates GSK3β on S9, which negatively regulates GSK3β kinase activity towards certain substrates. We next examined how GSK3β phosphorylation may contribute to control of GSK3β nuclear localization by mTORC1. As expected, cells treated with LY294002 or Akti-1/2 exhibited significant reductions in GSK3β S9 phosphorylation by 80 ± 0.8 % and 60 ± 6.8% respectively (n = 3, p < 0.05, **Fig. 5A**). In contrast, cells treated with rapamycin exhibited no change in GSK3β S9 phosphorylation compared to control (**Fig. 5A**). These results uncouple S9 phosphorylation from control of GSK3β nuclear localization. To directly probe the contribution of GSK3β S9 phosphorylation to mTORC1-dependent GSK3β nuclear localization, we studied the subcellular localization of GSK3β S9A. Under basal conditions, GSK3β S9A remains cytosolic, while treatment with the Akt inhibitor Akti-1/2 resulted in nuclear localization of GSK3β S9A, as seen with GSK3β WT (**Fig. 5B**).

**Figure 5.**
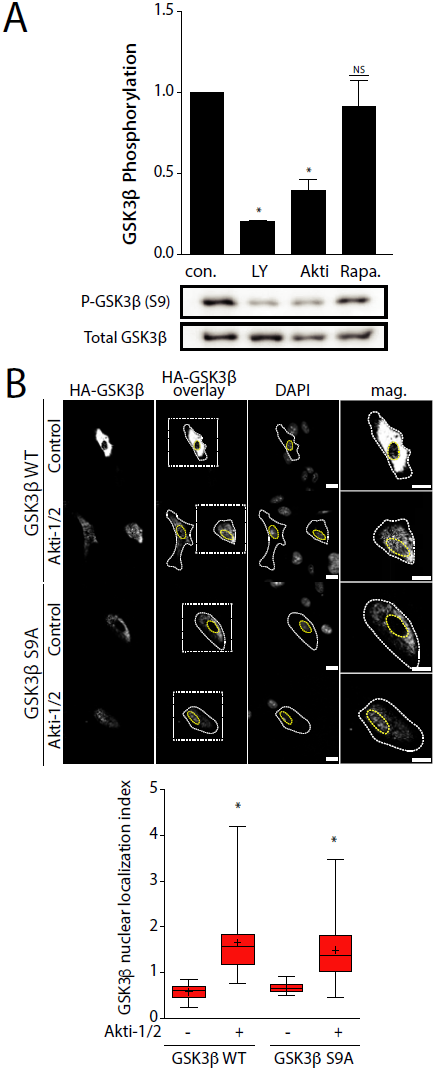
GSK3β S9 phosphorylation is not required for GSK3β nuclear localization induced by inhibition of PI3K-Akt-mTORC1 signals. (***A***) RPE cells were treated either 10 μM LY294002, 5 μM Akti-1/2, or 1 μM Rapamycin for 1 h. Shown are representative immunoblots of whole-cell lysates probed with anti-pS9 GSK3β or anti-total GSK3β antibodies. Also shown are mean anti-pS9 GSK3β levels (normalized to total GSK3β) ± SE, n = 3, * *p* < 0.05, relative to that in the control conditions (no inhibitor treatment). (***B***) RPE cells were transfected with plasmids encoding HA-tagged wild-type (WT) or S9A GSK3β then treated with 5 μM Akti-1/2 for 1 h, followed by detection of exogenous HA-GSK3β proteins. Shown (top panel) are micrographs obtained by widefield epifluorescence microscopy representative of 3 independent experiments, scale = 20 μm. Also shown for each condition as ‘HA-GSK3β overlay’ are sample cellular and nuclear outlines, and a box corresponding to a magnified image of a single cell. Also shown (bottom panel) is the mean HA-GSK3β nuclear localization index ± SE (n = 3, >30 cells per condition per experiment); *, *p* < 0.05 relative to control conditions (no inhibitor treatment).

Using phos-tag acrylamide electrophoresis, a technique that exaggerates differences in apparent molecular weight of phosphorylated species of a protein (Kinoshita et al., 2006), we observed two detectable species of GSK3β, of which the higher molecular weight species likely corresponds to the S9 phosphorylated form given its sensitivity to PI3K and Akt inhibition (**Fig. S4A**). In contrast and as expected, rapamycin had no effect on GSK3β detected by this method. Collectively, these results indicate that regulation of GSK3β S9 phosphorylation does not contribute to control of GSK3β nuclear localization by PI3K-Akt-mTORC1 signals.

### GSK3β is localized to several distinct membrane compartments within the cytoplasm

Active mTORC1 is recruited to the surface of the lysosome (Sancak et al., 2008). Together with our observations that mTORC1 controls GSK3β nuclear localization, this suggests that (i) mTORC1 control of GSK3β may occur at lysosomes and (ii) control of GSK3β nuclear localization by mTORC1 may require lysosomal membrane traffic. To determine if a pool of GSK3β is indeed localized to lysosomes concomitantly to GSK3β recruitment to other endomembrane compartments, we systematically examined the localization of endogenous GSK3β relative to APPL1 and EEA1 early endosomes, and to lysosomes demarked by LAMP1. We observed punctate distribution of endogenous GSK3β within the cytoplasm, with some visible overlap with each of APPL1, EEA1 and LAMP1 (**Fig. 6A-C**, left panels). To determine if the overlap observed between GSK3β and each marker was specific, we used quantification by Manders’ coefficient to compare overlap between real pairs of image channels, as well as between pairs of images with scrambled channel spatial position. This revealed specific GSK3β recruitment to each membrane compartment (**Fig. 6A-C**). We performed a similar colocalization analysis of the image data using Pearson’s coefficient and obtained similar results (**Fig. S4B**). This indicates that GSK3β indeed exhibits partial yet specific localization to several distinct endomembrane compartments, including APPL1 and EEA1 early endosomes, and late endosomes/lysosomes demarked by LAMP1.

**Figure 6.**
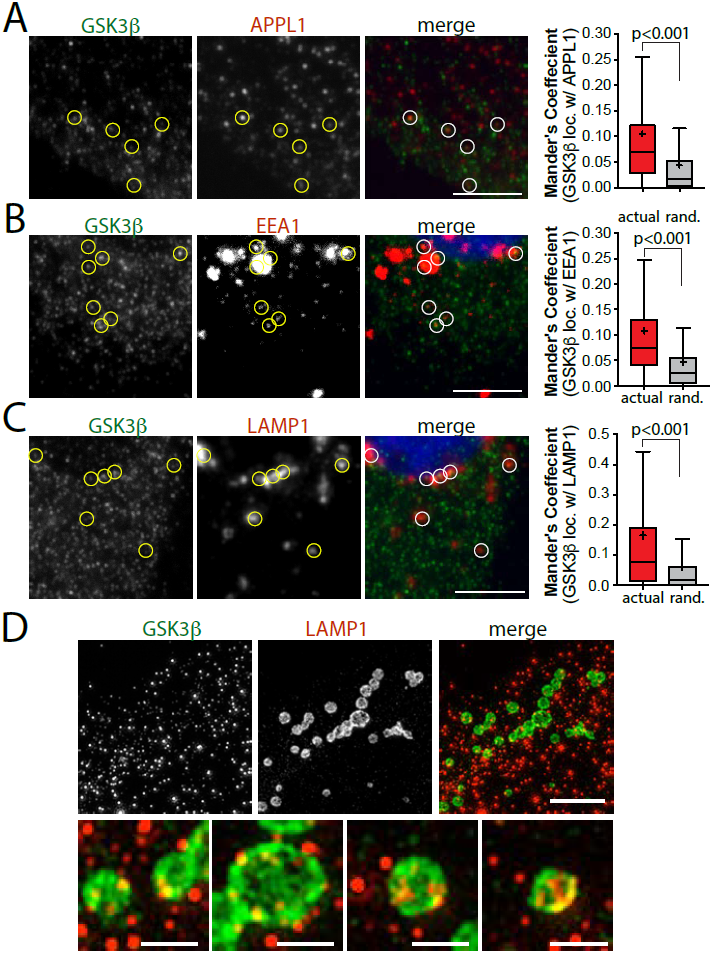
GSK3β exhibits partial localization to several distinct endomembrane compartments. (***A-C***) RPE cells were fixed and stained to detect endogenous GSK3β, together with either endogenous APPL1 (A), EEA1 (B), or LAMP1 (C). Shown are representative images obtained by spinning-disc confocal microscopy, corresponding to a z-section through the middle of the cell, scale 20 μm (left panels). Also shown (right panels) are the mean ± SE of Manders’ coefficient to measure overlap of GSK3β signals with either APPL1 (A), EEA1 (B), or LAMP1 (C) (n = 3, > 30 cells per condition per experiment). For each image set, Manders’ coefficients were calculated for actual images (labelled ‘actual’), as well as images in which the spatial position of one of the channels had been randomized (labelled ‘rand.’), to allow resolution of specific GSK3β localization to various endomembrane compartments from random overlap of signals in a field densely populated with fluorescent objects. (***D***) RPE cell samples prepared similarly as in (C) were subjected to structured illumination microscopy (SIM). Shown are representative micrographs of (endogenous) GSK3β and LAMP1 staining morphology, scale 5 μm (top panels) or 1 μm (bottom panel).

To further examine how GSK3β may localize to lysosomes, we employed structured illumination microscopy (SIM). Using this method, we were able to resolve the limiting membrane of lysosomes demarked by LAMP1 fluorescence staining (**Fig. 6D**). Importantly, GSK3β fluorescence staining was readily observed in punctate structures, in part associated with the limiting membrane of the lysosome. These results indicate that a subset of GSK3β in the cytoplasm exhibits association with the lysosome, either restricted to sub-domains of the lysosomal surface (Kaushik et al., 2006) or in structures associated with the lysosome, such as within membrane contact sites (Chu et al., 2015).

### Control of GSK3β nuclear localization and c-myc expression requires normal lysosomal membrane traffic

Given the localization mTORC1 (Sancak et al., 2008) and partial localization of GSK3β (**Fig. 6C-D**) to or near the lysosome, we next sought to determine the role of late endosome/lysosome membrane traffic to mTORC1-dependent control of GSK3β nuclear localization. To do so, we used a Rab7 mutant that is constitutively GDP-bound (T22N), which disrupts membrane traffic at the late endosome/lysosome (Choudhury et al., 2002). Cells expressing Rab7 T22N exhibited a significant increase in nuclear GSK3β, even in the absence of rapamycin treatment, compared to cells expressing Rab7 WT (**Fig. 7A**). Furthermore, cells expressing Rab7 T22N exhibited a depletion of GSK3β from lysosomes, observed by overlap of endogenous GSK3β and LAMP1, quantified by Manders’ coefficient (**Fig. S4C**). In contrast to the nuclear accumulation of GSK3β in cells expressing Rab7 T22N, silencing of APPL1 to disrupt early endosome membrane traffic did not impact GSK3β nuclear localization (**Fig. S4D**). These results indicate that membrane traffic at the late endosome/lysosome may be important to organize mTORC1 signals leading to control of GSK3β nuclear localization.

**Figure 7.**
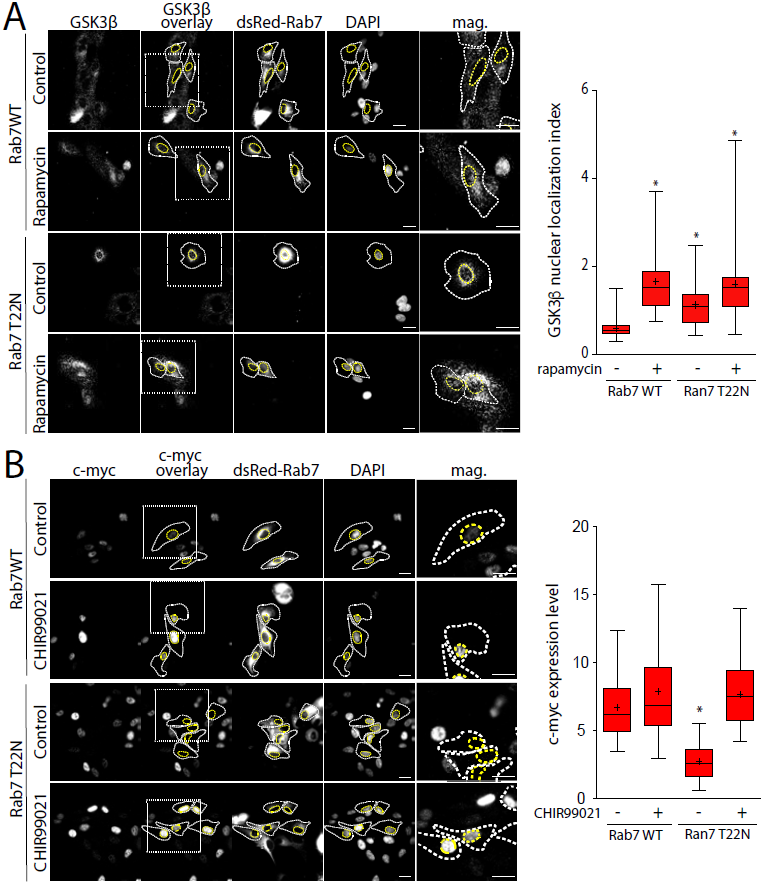
Rab7 controls GSK3β nuclear localization and GSK3β-dependent c-myc expression. RPE cells were transfected with plasmids encoding dsRed-tagged wild-type (WT) or T22N Rab7, then treated with 1 μM rapamycin for 1 h, followed by detection of endogenous GSK3β (***A***) or c-myc (***B***). Shown (left panels) are micrographs obtained by widefield epifluorescence microscopy representative of 3 independent experiments, scale = 20 μm. Also shown for each condition as ‘GSK3β overlay’ (A) or ‘c-myc overlay’ (B) are sample cellular and nuclear outlines, and a box corresponding to a magnified image of a single cell. Also shown (right panels) is the mean ± SE of the GSK3β nuclear localization index (A) (n = 3, >30 cells per condition per experiment) or total cellular c-myc level (B) (n = 3, >30 cells per condition per experiment); *, *p* < 0.05 relative to control conditions (no inhibitor treatment).

In order to determine the consequence of Rab7-dependent control of GSK3β nuclear localization, we examined the effect of expression of Rab7 T22N on GSK3β-dependent c-myc expression levels. Cells expressing Rab7 T22N exhibited a stark reduction in c-myc expression relative to cells expressing Rab7 WT (**Fig. 7B**). Importantly, treatment of cells expressing Rab7 T22N with the GSK3β inhibitor CHIR 99021 restored c-myc expression levels to that observed in cells expressing Rab7 WT (**Fig. 7B**). Taken together, these results indicate that control of GSK3β nuclear localization requires Rab7-dependent late endosome/lysosomal membrane traffic, reflecting perhaps the role of lysosomes as platform for mTORC1 signaling required to negatively regulate GSK3β nuclear translocation.

## Discussion

We identified that the nuclear localization of GSK3β is regulated by mTORC1, such that conditions that reduce mTORC1 activity result in increased nuclear localization of GSK3β, and increased GSK3β-dependent degradation of nuclear substrates such as c-myc and snail. Furthermore, GSK3β was partly but specifically localized to the surface of late endosomes/lysosomes, and perturbation of membrane traffic at the late endosomes/lysosomes disrupted GSK3β nucleocytoplasmic shuttling and regulation of c-myc expression.

### Localization of GSK3β within multiple membrane compartments within the cytoplasm

Separate studies have reported that GSK3β may localize to a number of distinct cellular compartments, including endomembranes, mitochondria and the nucleus (reviewed by (Beurel et al., 2015)). By a systematic, unbiased approach, we find that endogenous GSK3β localizes to several distinct endomembrane compartments, including APPL1 endosomes, EEA1-positive early endosomes and LAMP1-positive late endosomes/lysosomes (**Fig. 6**). In each case, the overlap of GSK3β immunofluorescence signal and that of each compartment marker is clearly limited and partial, with substantial proportions of each signal not exhibiting overlap (**Fig. 6A-C**). However, systematic and unbiased analysis of colocalization performed by Manders’ (**Fig. 6A-C**) or Pearson’s (**Fig. S4B**) coefficient analysis indicates that GSK3β overlap with each compartment is specific and non-random. Moreover, the specific recruitment of GSK3β to the limiting membrane of LAMP1-positive late endosomes/lysosomes is supported by images obtained by SIM (**Fig. 6D**), as well as by the observation that perturbation of late endosome/lysosome membrane traffic by expression of a dominant interfering mutant of Rab7 abolishes the overlap of GSK3β with LAMP1 signals (**Fig. S4C**). Our observations are thus consistent with the notion that GSK3β is localized to a number of distinct cellular compartments, with a minor pool that in some cases is <10% of total cellular GSK3β, recruited to each such compartment at steady state.

Our observations are also consistent with previous studies showing GSK3β localization to APPL1 endosomes (Schenck et al., 2008). APPL1 recruitment to a subset of internalized membranes formed by clathrin-mediated endocytosis precedes the acquisition of markers of the EEA1 early endosome (Zoncu et al., 2009). This pool of GSK3β within APPL1 endosomes may be specifically targeted by phosphorylation on S9 by Akt, as silencing of APPL1 abolishes Akt-dependent GSK3β phosphorylation (Reis et al., 2015; Schenck et al., 2008). Notably, perturbation of APPL1 by silencing did not impact mTORC1-dependent control of GSK3β nuclear localization (**Fig. S4D**), suggesting that the APPL1-localized pool of GSK3β does not directly participate in the regulation of GSK3β nuclear localization by mTORC1.

As mTORC1 localizes to the surface of late endosomes and lysosomes, the pool of GSK3β on these membranes may be under the direct regulation by mTORC1 to control GSK3β nucleocytoplasmic shuttling. Indeed a previous report had observed some overlap of GSK3β and the lysosome (Li et al., 2016). However, GSK3β may also be sequestered within intraluminal vesicles of multivesicular bodies in response to Wnt signaling (Taelman et al., 2010), raising the possibility that the overlap that we observed by spinning disc confocal microscopy between LAMP1-positive structures and GSK3β (**Fig. 6C**) could reflect GSK3β within intraluminal vesicles. However, examination of SIM images suggests that very little, if any, GSK3β is observed within the lumen of LAMP1-positive structures (**Fig. 6D**), suggesting that LAMP1-localized GSK3β is largely associated with the limiting membrane of these compartments. Moreover, perturbation of Rab7 disrupts the localization of GSK3β and LAMP1 (**Fig. S4C**), yet Rab7 disruption does not impact the sequestration of material into intraluminal vesicles (Vanlandingham and Ceresa, 2009). The molecular mechanism(s) by which GSK3β is recruited to lysosomes remains unknown, and is beyond the scope of this study. Our results thus add systematic analysis and quantification to indicate that a pool of GSK3β is present on the limiting membrane of the lysosome, and suggesting that this pool may be subject to regulation by mTORC1, resulting in control of GSK3β nuclear localization.

### Mechanism of control of GSK3β nuclear localization by mTORC1

We found that direct inhibition of any component of the PI3K-Akt-mTORC1 axis, or activation of AMPK to trigger mTORC1 inhibition results in an increase in GSK3β nuclear localization. Moreover, perturbation of Rab7-dependent membrane traffic also resulted in an increase in GSK3β nuclear localization, suggesting that in addition to mTORC1 signals, lysosomal traffic and/or organization is also required to control GSK3β nuclear import. Interestingly, we also observed that inhibition of PI3K-Akt-mTORC1 also increased nuclear localization of GSK3α. Hence, it is likely that mTORC1 signals similarly gate GSK3α and GSK3β nuclear localization. Taken together, we propose that mTORC1 establishes a form of ‘molecular licencing’ for retention within the cytoplasm for GSK3α and GSK3β, resulting in nuclear exclusion under conditions of elevated mTORC1 activity. This molecular licencing could take the form of a post-translational modification of GSK3α or GSK3β, or of regulation of protein complex formation at specific subcellular locale(s).

GSK3β undergoes nucleocytoplasmic shuttling, due to nuclear import in balance with FRAT-1-mediated nuclear export (Wiechens and Fagotto, 2001). Nuclear import of some (but not all) proteins is controlled by a gradient of GTP-bound and GDP-bound Ran that spans the nuclear membrane (Strambio-De-Castillia et al., 2010). By expression of mutants of Ran (**Fig. 3**), we show that the nucleocytoplasmic shuttling of GSK3β is Ran-dependent. Nuclear import of GSK3β resulting from mTORC1 inhibition by rapamycin was prevented in cells expressing Ran G19V mutant defective in GTP hydrolysis and thus defective in nuclear import. Hence, nuclear import of GSK3β regulated by mTORC1 is Ran-dependent.

We examined whether the phosphorylation of S9 on GSK3β could control its mTORC1-regulated nuclear localization; however, two observations strongly suggest that this is not the case: (i) inhibition of mTORC1 by rapamycin did not alter S9 phosphorylation of GSK3β (**Fig. 5A**), and (ii) a mutant of GSK3β that cannot be phosphorylated at this position (S9A) behaved similarly to wild-type with respect to mTORC1-dependent nuclear localization (**Fig. 5B**). GSK3β can also be phosphorylated on a number of other residues, including Y216, which may result from auto-phosphorylation at the time of GSK3β synthesis (Beurel et al., 2015). Further, GSK3β can be phosphorylated at T43 (Ding et al., 2005) and S389 (Thornton et al., 2008) by Erk and p38 MAPK, respectively, each of which lead to reduction in GSK3β activity. Notably, using a phos-tag gel electrophoresis approach, a technique that exacerbates the apparent molecular weight increase caused by phosphorylation, we were only able to resolve two bands for GSK3β that likely correspond to S9 phosphorylated and non-S9 phosphorylated forms (**Fig. S4A**). It will be interesting to determine in future studies if and how phosphorylation at sites other than S9 are regulated by mTORC1 to control GSK3β nuclear localization.

Other than phosphorylation, other modifications reported for GSK3β include citrullination (Stadler et al., 2013) ADP-ribosylation (Feijs et al., 2013) and calpain cleavage (Goñi-Oliver et al., 2007). Indeed, citrullination of R3 and R5 residues within GSK3β is important for nuclear localization (Stadler et al., 2013). However, we observed that mTORC1 controls both GSK3α and GSK3β nuclear localization, and these two GSK3 paralogs differ at their N-terminus within the region of GSK3β that undergoes citrullination. Hence, it appears unlikely to expect that mTORC1 controls citrullination of GSK3β as a mechanism of control of its nucleocytoplasmic shuttling. While beyond the scope of this study, it will be interesting to note how future work may resolve whether mTORC1-dependent regulation of post-translational modification of GSK3β underlies the regulation of its nuclear localization by mTORC1.

mTORC1-dependent control of GSK3β nuclear localization may occur as a result of regulation of GSK3β interaction with other proteins in various endomembrane compartments. It is worth noting that the vast majority of cytoplasmic, but not nuclear GSK3β, is associated with other protein(s) (Meares and Jope, 2007). Thus, it is possible that control of GSK3β nucleocytoplasmic shuttling involves regulation of protein-protein interactions that serve to occlude the bipartite NLS of GSK3β (residues 85 to 103) (Meares and Jope, 2007), thus limiting GSK3β nuclear localization when these interactions are present.

We also found that Rab7 is required to retain GSK3β in the cytoplasm under conditions when mTORC1 is otherwise active. Importantly, disruption of late endosome/lysosome membrane traffic by perturbations of Rab7 or other proteins does not impact mTORC1 activity (Flinn et al., 2010). This indicates that the ability of mTORC1 to limit the nuclear localization of GSK3β requires active traffic to the late endosome/lysosome. This in turn suggests that the protein interactions engaged by GSK3β that occlude its NLS and thus limit nuclear localization may occur on the lysosome, consistent with our observed localization of GSK3β to the lysosome. Indeed GSK3α and GSK3β have nearly identical kinase domains (in which the NLS is found), consistent with the ability of mTORC1 to gate nuclear access for both GSK3 paralogs.

Furthermore, our observations that mTORC1 controls GSK3β nuclear localization add to previous reports that GSK3β activates mTORC1 signals (Inoki et al., 2006), and suggests the existence of reciprocal regulation of mTORC1 and GSK3β. Overall, we propose that mTORC1 signals limit the ability of GSK3β to localize to the nucleus, and that this may result from mTORC1-dependent control of GSK3β interactions with other proteins in a manner that regulates occlusion of the NLS of GSK3β at the lysosome.

### Regulation of GSK3 β nuclear functions by mTORC1

We identified that various metabolic and mitogenic signals gate nuclear access for GSK3β. This in turn allows for GSK3β-dependent regulation of nuclear substrates in response to mTORC1 signals. Previous studies reported that nuclear and cytoplasmic pools of GSK3β have distinct functions, such as nuclear GSK3β facilitating stem cell differentiation over self-renewal (Bechard and Dalton, 2009) or the cytosolic pool of GSK3β being sufficient to mediate GSK3β-dependent cell survival to tumor necrosis factor α (TFNα) apoptotic signals (Meares and Jope, 2007).

One of the nuclear substrates of GSK3β is c-myc, a helix-loop-helix-leucine zipper transcription factor that has a very short half-life (15-30 mins) (Kalkat et al., 2017; Lüscher and Eisenman, 1988). As previously reported, nuclear localization of GSK3β is required for phosphorylation of GSK3β on T58, resulting in enhanced c-myc degradation (Gregory et al., 2003). We show that rapamycin treatment, which promotes nuclear localization of GSK3β, also results in an acute reduction in c-myc accumulation (**Fig. 1**), most likely due to c-myc degradation. A previous report suggested that rapamycin treatment elicits degradation of c-myc by induction of autophagy, as result of induction of AMBRA-dependent dephosphorylation of c-myc at T58 (Cianfanelli et al., 2014). However, we show that the degradation of c-myc induced by rapamycin is insensitive to impairment of autophagy induction elicited by siRNA gene silencing of ULK1 (**Fig. S3**). Moreover, we find that the rapamycin-induced reduction in c-myc levels is countered by perturbation of GSK3β (**Fig. 1A-B**). Hence, our results indicate that mTORC1-dependent control of GSK3β nuclear localization regulates c-myc in a manner that does not require induction of autophagy.

Based on the control of GSK3β nuclear localization by mTORC1 leading to control of c-myc, we propose the existence of a metabolic sensing signaling network that links nutrient availability with biomass production and proliferation. Indeed, c-myc controls the expression of many genes, generally to promote ribosome production, biomass accumulation and enhanced cellular bioenergetics, such as through mitochondrial biosynthesis (Miller et al., 2012). Furthermore, c-myc promotes epithelial-mesenchymal transition (Cho et al., 2010). Hence, signals activated during nutrient deficiency can impair the anabolic c-myc-dependent promotion of biomass accumulation via this novel mTORC1-GSK3β-c-myc signaling axis involving control of GSK3β nuclear localization.

GSK3β may also regulate other nuclear substrates selectively during conditions of reduced mTORC1 signaling or other states in which GSK3β exhibits nuclear localization. Collectively, regulation of other GSK3β substrates such as snail (leading to degradation, (Sekiya and Suzuki, 2011)) or c-jun (leading to impaired DNA binding, (Nikolakaki et al., 1993)) is consistent with the effect of GSK3β-dependent degradation of c-myc: reduced cell cycle progression, impairment of epithelial-mesenchymal transition and/or reduced biomass accumulation. While examination of mTORC1-dependent regulation of all known GSK3β nuclear targets is beyond the scope of this study, it is perhaps tempting to speculate that metabolic and mitogenic signals broadly control the nuclear profile of GSK3β functions, coordinating energy-demanding accumulation of biomass, cell cycle progression and growth with nutrient availability. As cancer cells exhibit heterogeneity of metabolic cues and signals, it is possible that differences in metabolism between cancer cells that result in distinct GSK3β nuclear localization profiles may underlie in part the differences in response to drugs targeting GSK3β in cancer, although this remains to be examined.

In conclusion, we identified that GSK3β nucleocytoplasmic shuttling is controlled by both mitogenic signals such as PI3K-Akt and metabolic cues including amino acid or ATP availability as a result of mTORC1-dependent control of GSK3β nuclear import. In addition, GSK3β localized in part to the late endosome/lysosome and nuclear localization of GSK3β was regulated by Rab7, suggesting that membrane traffic at late endosomes and lysosomes impacts signals leading to control of GSK3β nuclear localization. Lastly, we propose that GSK3β-dependent control of nuclear proteins by mTORC1 occurs by regulation of GSK3β nuclear import, linking nutrient availability to control of energy-dependent transcriptional networks.

## Materials and Methods

### Materials

Antibodies targeting specific proteins were obtained as follows: GSK3β, phospho-GSK3β (S9) actin, HA-epitope, EEA1, LAMP1, APPL1, and ULK1 (Cell Signaling, Danvers, MA), and clathrin (Santa Cruz Biotechnology, Dallas, TX). Horseradish peroxidase or fluorescently-conjugated conjugated secondary antibodies were purchased from Cell Signaling Technology (Danvers, MA). DAPI Nuclear staining was purchased from ThermoFisher (Rockford, IL).

Ran cDNA constructs tagged to HA, including WT, T24N and G19V forms in pKH3 were generously provided by Dr. Ian Macara (Vanderbilt University School of Medicine, Nashville, TN) (Carey et al., 1996). GSK3β cDNA constructs, including HA-tagged WT and S9A forms in pcDNA3 were generously provided by Dr. Jim Woodgett (Lunenfeld-Tanenbaum Research Institute/Mount Sinai Hospital, Toronto, ON) (Stambolic and Woodgett, 1994). Rab7 constructs, including WT and T22N, were generously provided by Dr. Richard Pagano (Mayo Clinic and Foundation, Rochester, MN) (Choudhury et al., 2002).

### Cell lines, cell culture and inhibitor treatment

Wild-type human retinal pigment epithelial cells (ARPE-19; RPE herein) were cultured were obtained from American Type Culture Collection (ATCC, Manassas, VA) as previously described (Delos Santos et al., 2017) with DMEM/F12 (Gibco, ThermoFisher Scientific, Waltham, MA) containing 10 % fetal bovine serum, 100 U/ml penicillin and 100 μg/ml streptomycin. Cells were then incubated at 37 C and 5 % CO_2_. MDA-MB-231 cells were obtained from ATCC and cultured as previously described (Fekri et al., 2016) with RPMI media 1640 (Gibco) containing 10 % fetal bovine serum, 100 U/ml penicillin and 100 μg/ml streptomycin and incubated at 37C and 5 % CO_2_.

All inhibitor treatments were performed (alone or in combination) for 1 h prior to experimental assays unless otherwise indicated, as follows: 10 μM CHIR 99021 (Abcam, Cambridge, MA), 1 μM Rapamycin (BioShop, Burlington, ON), 10 μM LY294002 (Cell Signaling Technologies), 5 μM Akti-1/2 (Sigma-Aldrich, Oakville, Canada), 1 μM Concanamycin A (BioShop). Amino acid starvation was performed by incubation in Earle’s Balanced Salt Solution (EBSS, Gibco).

### Plasmid and siRNA transfections

To perform DNA plasmid transfections, Lipofectamine 2000 (ThermoFisher Scientific) was used according to the manufacturers instructions and as previous described (Bone et al., 2017). Briefly, cells were incubated for 4 h with Lipofectamine 2000 reagent and appropriate plasmid in Opti-MEM (Gibco) at a 3:1 ratio. Subsequently, this transfection solution was removed, and cells were incubated in fresh cell growth medium at 37C and 5% CO_2_ for 16-24 h prior to experimentation.

To perform siRNA transfections as previously described (Bone et al., 2017), custom-synthesized siRNAs targeting specific transcripts with sequences as follows were obtained from Dharmacon (Lafayette, CO) as follows: non-targeting control: CGU ACU GCU UGC GAU ACG GUU (sense strand), and CGT ACT GCT TGC GAT ACG GUU (antisense strand); GSK3β: ACA CUA UAG UCG AGC CAA AUU (sense strand), and UUU GGC UCG ACU AUA GUG U (antisense strand); ULK1: GCA CAG AGA CCG UGG GCA AUU (sense strand), and UUG CCC ACG GUC UCU GUG CUU (antisense strand); APPL1: CAG AAU GUU CGC AGG GAA AUU (sense strand), and UUU CCC UGC GAA CAU UCU GUU (antisense strand). Cells were incubated with 220 pmol/L of each siRNA sequence with Lipofectamine RNAiMAX (LifeTechnologies) in Opti-MEM medium (Gibco) for 4 hours at 37C and 5% CO_2_. After this incubation period, cells were washed and incubated in fresh cell growth medium. siRNA transfections were performed twice, 72h and 48h prior to experiments.

### Whole-cell lysates and Western blotting

Western blotting using whole-cell lysates were performed as previously described (Garay et al., 2015). Cells were lysed in Laemmli sample buffer (LSB; 0.5 M Tris, pH 6.8, glycerol, 5% bromophenol blue, 10% β-mercaptoethanol, 10% SDS; BioShop, Burlington, ON) containing phosphatase and protease cocktail (1 mM sodium orthovanadate, 10 nM okadaic acid, and 20 nM protease inhibitor, all from BioShop, Burlington, ON). Cell Lysates were then heated to 65C for 15 min, then passed through with a 27.5-gauge syringe. Proteins within whole-cell lysates were resolved by Glycine-Tris SDS-PAGE and then transferred onto a polyvinylidene fluoride (PVDF) membrane, which was then incubated with a solution containing specific primary antibodies. Western blot signal intensity detection corresponding to either phosphorylated proteins (e.g. pGSK3β S9), total proteins (e.g. GSK3β), and the respective loading controls (e.g. actin) were obtained by signal integration in an area corresponding to the specific lane and band for each condition. The measurement of phosphorylation of a specific protein was obtained by normalization of the signal intensity of a phosphorylated form a protein to that of its loading control signal, then normalization to the signal intensity similarly obtained for the corresponding total protein.

To examine phosphorylation of proteins for which no specific antibodies were available, we used the phos-tag gel system, which results in exaggeration of differences in apparent molecular weight of phosphorylated forms of specific proteins (Kinoshita et al., 2006). The phos-tag reagent was obtained from Wako (Osaka, Japan), and was used for conjugation within SDS-PAGE polymerization as per the manufacturer’s instructions. After SDS-PAGE was completed, gel was submerged in MnCl_2_ for chelation of remaining phos-tag moieties. Subsequently, protein intensity detection, measurement, and processing are identical to steps mentioned above.

### Immunofluorescence staining

Cells grown on glass coverslips were first subjected to fixation using cold methanol, blocked in 5% bovine serum albumin (BioShop), then stained with specific primary antibodies, followed by appropriate fluorophore-conjugated secondary antibody and counter stained with DAPI. Lastly, cells were then mounted on glass slides in fluorescence mounting medium (DAKO, Carpinteria, CA).

### Fluorescence microscopy

Wide-field epifluorescence was performed on an Olympus IX83 Inverted Microscope with a 100x objective, coupled to to a Hamamatsu ORCA-Flash4.0 digital camera (Olympus Canada, Richmond Hill, ON). Spinning disk confocal microscopy was performed using Quorum (Guelph, ON, Canada) Diskovery combination total internal reflection fluorescence and spinning-disc confocal microscope, operating in spinning disc confocal mode. This instrument is comprised of a Leica DMi8 microscope equipped with a 63×/1.49 NA objective with a 1.8× camera relay (total magnification 108×). Imaging was done using 488-, 561-, and 637-nm laser illumination and 527/30, 630/75, and 700/75 emission filters and acquired using a Zyla 4.2Plus sCMOS camera (Hamamatsu).

Structured illumination microscopy (SIM) was performed using a Zeiss Elyra PS.1 super-resolution inverted microscope, as previously described (Hua et al., 2017). Samples were imaged at an effective magnification of 101x (63x objective + 1.6x optovar tube lens) on an oil immersion objective. Typically, 25 to 35 slices of 0.110 μm were captured for each field of view for an imaging volume of approximately 2.75 to 3.85 μm. 488 nm, 561 nm and 643 nm laser lines were directed into the microscope optical train via a multimode fiber coupler. The lasers were passed through a diffraction grating, and a series of diffraction orders (-1, 0, +1) were projected onto the back focal plane of the objective. These wavefronts were collimated in the objective to create a three-dimensional sinusoidal illumination pattern on the sample. The diffraction grating was then rotated and translated throughout the acquisition to create patterned offset images containing encoded high spatial frequency information. Five lateral positions were acquired at each of five (72°) diffraction grating rotations for a total of 25 raw images per z-plane. SIM imaging with all lasers was carried out at exposures varying from 50 ms to 100 ms, with laser power varying between 3-10% (6-20 mW at the output), and a gain level of 60-80. Imaging parameters were adjusted iteratively to achieve the best possible equalization of pixel intensity dynamic range across channels.

Raw SIM image stacks were processed in Zen under the Structured Illumination toolbar. A series of parameters were set to generate an optical transfer function (OTF) used for 3D reconstruction. The noise filter for Wiener de-convolution was set to a value of 1.0 × 10^−4^ to maximize the recovery of high spatial frequency information while minimizing illumination pattern artifacts. The maximum isotropy option was left unselected to recover all available frequency information at exactly the 72° rotation angles. Negative values arising as an artifact of the Wiener filter were clipped to zero using the Baseline Cut option. Processed SIM images were then aligned via an affine transformation matrix of pre-defined values obtained using 100 nm multicolor Tetraspeck fluorescent microspheres (ThermoFisher Scientific).

### Fluorescence microscopy image analysis

Measurement of total cellular signal intensity of specific proteins or GSK3β nuclear localization index were measured using ImageJ software (National Institutes of Health, Bethesda, MD). For total cellular measurements of specific protein signal, a region of interest corresponding to the cell outline, identified manually, was used to determine raw mean cellular fluorescence intensity. Final cellular signal intensity was obtained by subtracting background fluorescence (similarly obtained from a region on the coverslip with no cells) from the raw mean cellular fluorescence intensity, as previously described (Ross et al., 2015).

To determine GSK3β nuclear localization index, background-subtracted mean fluorescence intensity of regions of interest within the nucleus and cytoplasm were obtained. The GSK3β nuclear localization index is the ratio of these nuclear/cytosolic intensities. Each measurement was performed in at least three independent experiments, with > 30 cells per condition, per experiment.

Colocalization analysis was performed by determination of Manders’ or Pearson’s coefficients, as indicated, using the Just Another Colocalization Plugin (Bolte and Cordelières, 2006) within ImageJ, as previously described (Bone et al., 2017).

### Statistical analysis

Statistical analysis was performed as previously described (Bone et al., 2017). Measurement of samples involving two experimental conditions (**Figs. 3B, 4B, 6, S2, S3A & S4B**) were analyzed by student’s t-test, with p < 0.05 as a threshold for statistically significant difference between conditions. Measurements of samples involving one experimental parameter and more than two conditions (**Figs. 1, 2B, 4A, 4C & S1**) were analyzed by one-way ANOVA, followed by Bonferonni post-test to compare differences between conditions, with p < 0.05 as a threshold for statistically significant difference between conditions. Measurements of samples involving two experimental parameters (**Figures 3, 5, 7, S3B, S3C, S4C & S4D**) were analyzed by two-way ANOVA, followed by Bonferonni post-test to compare differences between conditions, with p < 0.05 as a threshold for statistically significant difference between conditions.

## Acknowledgements

We thank Dr. Jim Woodgett (Lunenfeld-Tanenbaum Research Institute/Mount Sinai Hospital, Toronto, ON) and Dr. Sean Egan (Hospital for Sick Children, Toronto, ON) for insightful and helpful discussions. This work was supported by a Discovery Grant from the Natural Sciences and Engineering Research Council (of Canada), an Early Researcher Award from the Ontario Ministry of Research, Innovation and Science, and a New Investigator Award from the Canadian Institutes of Health Research to C.N.A.

